# Urological benchtop and *in silico* models validated by human penile tissue inflation tests

**DOI:** 10.1101/2025.06.18.660316

**Authors:** Majid Akbarzadeh Khorshidi, Shirsha Bose, Brian Watschke, Evania Mareena, Thomas Sinnott, Caitríona Lally

## Abstract

**Purpose:** Inflatable penile prosthesis (IPP) implantation is a well-established treatment for erectile dysfunction (ED). A comprehensive understanding of the mechanical interactions between the IPP and penile tissues is crucial for improving surgical outcomes and device performance. This study aims to develop and validate preclinical testbeds, including a polymer-based benchtop model and a finite element (FE) model, to replicate the biomechanical behaviour of penile tissues during IPP inflation.

**Methods:** A polymer-based benchtop model was developed using porous and non-porous polyvinyl alcohol (PVA) hydrogels, with the porous PVA mimicking the spongy corpus cavernosum (CC) and the non-porous PVA representing the tunica albuginea (TA) and fascial layers. IPP inflation tests were conducted on three benchtop models and three human penile tissue segments. Additionally, 3D FE simulations of IPP inflation were performed on both the benchtop and human tissue models for comparative analysis.

**Results:** The experimental results demonstrated strong agreement between the human penile tissues, the benchtop model, and the FE simulations, validating the preclinical testbeds. Parametric studies using the FE model revealed that CC layer size and stiffness significantly influence IPP inflation mechanics, highlighting the importance of these factors in device performance. These validated preclinical testbeds provide a robust platform for optimising IPP design, guiding surgical procedures, and mitigating post-implantation complications associated.

**Conclusion:** The developed benchtop and FE models effectively replicate human penile tissue responses to IPP inflation and can serve as valuable preclinical tools for device manufacturers and clinicians. Their use may enhance surgical decision-making and improve long-term IPP outcomes.

## 1. Introduction

Erectile dysfunction (ED) is a commonly under treated male sexual disease affecting the older male population, occurring due to the inability to attain or maintain erection. Several treatment methods for example, pharmacotherapy, vacuum erection devices, extracavernous injection etc. exist, however, patients with previous records of pelvic trauma, diabetes, vascular diseases might not respond to these conventional treatments and are required to undergo penile implantation via surgery [1]. Inflatable penile prostheses (IPPs) are one of the most common commercially available penile implants. IPPs consist of a pair of cylinders which is placed in the corpus cavernosa of the human penis facilitating in the erection of the penis. IPPs also contain a pump (in the scrotal sacs) and a reservoir (in the abdomen) [2].

Although the first pair of penile prosthesis was developed in 1952 by Goodwin and Scott [3], understanding the impact of these implants on the neighbouring tissues is not well explored yet. Some studies investigated the mechanics of penile tissue [4]–[11] and IPP [12]–[14], but further research is needed to predict the stress and deformation in the tissue caused by device inflation. Successful medical device implantation also requires strong understanding of the tissue biomechanics and development of suitable pre-clinical testbeds [15]. Validated computational and synthetic (benchtop) models will strongly aid to enhance our understanding of pre- and post-implantation tissue mechanical behaviour and tissue-device interactions, such as tissue damage after IPP inflation. This also helps clinicians to diminish the implantation complications by choosing the best implantation scenario [4]. On the other hand, these pre-clinical testbeds can contribute to device design and manufacturing process to achieve an optimal design.

Polyvinyl alcohol (PVA) is a biocompatible polymer with great potential for biomedical applications ranging from living organisms and function in disease treatment, physiological signal monitoring [16]–[18], scaffolds for tissue engineering and drug-delivery vehicles [19], [20]. In addition, PVA has found extensive use in replicating biological tissue behaviour [21]–[26]. Moreover, he mechanical properties of PVA can be tuned using different techniques, such as freeze-thaw-cycles (FTCs), PVA concentration, freezing temperature, etc. [19], [26]–[28]. This capability facilitates achieving the desired PVA properties that align with those of natural tissues. When considering tissue-mimicking applications, it is crucial not only that synthetic materials have convenient properties but also that they accurately replicate the mechanical responses of actual tissue or organs under physiological or supra-physiological loads, such as those imposed by medical devices. PVA is a synthetic polymer widely used for its non-toxicity, excellent mechanical properties and ability to tune its properties via physical or chemical cross-linking agents [29]. Physical cross-linking is achieved generally through freeze-thaw cycles [30] while chemical cross-linking can be achieved using various agents such as glutaraldehyde, boric acid, glyoxal etc. [31], [32]. Our research team has recently developed novel macro-porous PVA hydrogels [33] with potential applicability in soft spongy tissues like the corpus cavernosum (CC) in penile tissue. Since CC compresses as the IPP cylinder inflates within the tissue, this porous material can be compressed more during the IPP inflation compared to non-porous materials.

The aim of this study is to develop a validated penile benchtop model using porous PVA to replicate the CC layer and non-porous PVA to mimic the fascia-tunica layers. In addition, a 3D FE model of penile tissue is developed to simulate the IPP inflation within the tissue. An experimental set-up is designed to test both human penile tissue and the benchtop model under the IPP device inflation.

## 2. Materials and Methods

### 2.1. Human Tissue Preparation

Fresh human penises (N=3, with a post-mortem time interval of 1 – 5 days) were purchased from Science Care, USA with donors having age range from 75 – 89 years. The specimens were cleaned (Fig. 1a) with a scalpel to remove the unwanted neighbouring tissues. This was followed by removal of the glans of the penis (Fig. 1b) and then sectioned to 30 mm segments (n=3) using a custom-made cutter and microtome blades (Fig. 1c). These tissues were snap frozen with liquid nitrogen and stored in a freezer at – 20 °C until the test.

**Fig. 1:**
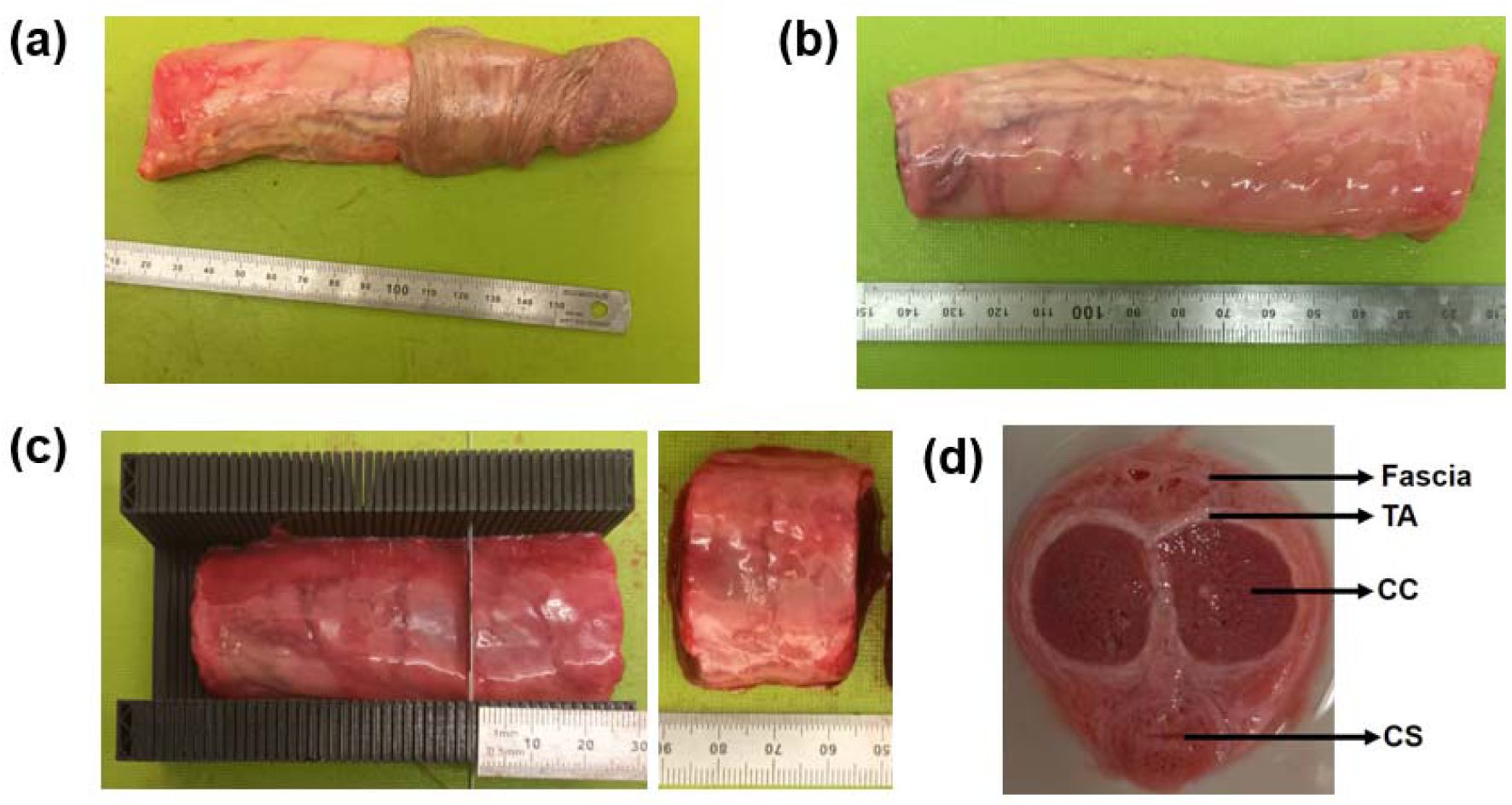
(a) Human penis with glans intact, (b) Removal of glans, (c) Custom-made cutter used to form 30 mm segments and (d) Cross-sectional view of the penile anatomy.

For defrosting, the tissues were placed in a pre-warmed water bath at 37 °C followed by removal of residual blood from the tissue by cleaning them with phosphate-buffered saline (PBS). The cross-sectional anatomy of the human penis is presented in Fig. 1d showing the pair of spongy erectile tissue chambers called CC Encompassing the CC, is a thin fibrous layer called TA which is further surrounded by fascial layers. Other penile components include the urethra which is enclosed by more spongy tissue called corpus spongiosum (CS).

### 2.2. Materials

The PVA powder used to prepare PVA solution (= 89,000-98,000 g/mol, >99% hydrolysed) was purchased from Sigma-Aldrich (Sigma-Aldrich, USA). Other ingredients used were aluminium oxide (544833-50G, Sigma Aldrich, UK and Merck) nano-powder (particle size <50 nm), glycerol (M_W_= 92.10 g/mol, Sigma Aldrich Germany) and 3D printed PVA filament (Ultimaker PVA Natural, 750g, Inspire 3D) . All the materials purchased were used without further modifications.

### 2.3. Preparation of PVA hydrogels

The preparation of both macro-porous and non-porous PVA hydrogels was achieved by following the protocol described by Bose et al. [33]. In brief, water-soluble sacrificial milled PVA (m-PVA), with particle sizes 300 – 500 µm, was obtained by milling 3D printed PVA particles in a cryo-miller operated at room temperature.

PVA solutions (5% w/v and 10% w/v) were prepared using PVA powder and other ingredients aluminium oxide (1.45%), glycerol (5%) and de-ionised (DI) water. The mixture was placed in a warm water bath 80 – 85 °C with continuous stirring with a magnetic stirrer for an hour to obtain a uniform and homogenous solution. The solution was cooled to room temperature before pouring it into moulds for non-porous PVA. For porous PVA, 35% (w/v) of sacrificial fillers were added to 5% PVA solution and slightly stirred and transferred to the required moulds. The moulds were then placed in a refrigerator at - 20°C for 14 h. Following, freezing it underwent thawing for 10 h. Upon completion of the required number of FTCs, the porous PVA specimen was placed in DI water for particulate leaching of the soluble m-PVA for 72 h. The PVA hydrogels were always stored in DI water at room temperature until testing (within 14 days of specimen preparation). Previous studies on degradation of PVA-based composites showed slight changes in mechanical properties after two weeks of preparation [34].

### 2.4. Development of 30 mm benchtop moulds

The geometry of the 30 mm segmented penis was developed in Abaqus/CAE (Simulia) and then exported to SOLIDWORKS^®^ 3D mechanical CAD software (version 2021, Dassault Systems) to create moulds for the benchtop model. The multi-layered penile anatomy (Fig. 2a) was simplified to have the CC and the neighbouring fascia-TA layers (Fig. 2b). This simplification was done as the IPP device during its inflation and deflation mostly affects the CC tissues. The mould was designed by extruding ‘Part A’ (Fig. 2c) to 40 mm and the fascia-TA layers to 10 mm. The CC was extruded to 5 mm. A pair of mandrels having the same geometry as the CC were also created as shown in Fig. 2d. Following the design of the moulds in SOLIDWORKS, they were printed in Original Prusa MK4 3D printer using Polylactic acid (PLA) filament.

**Fig. 2:**
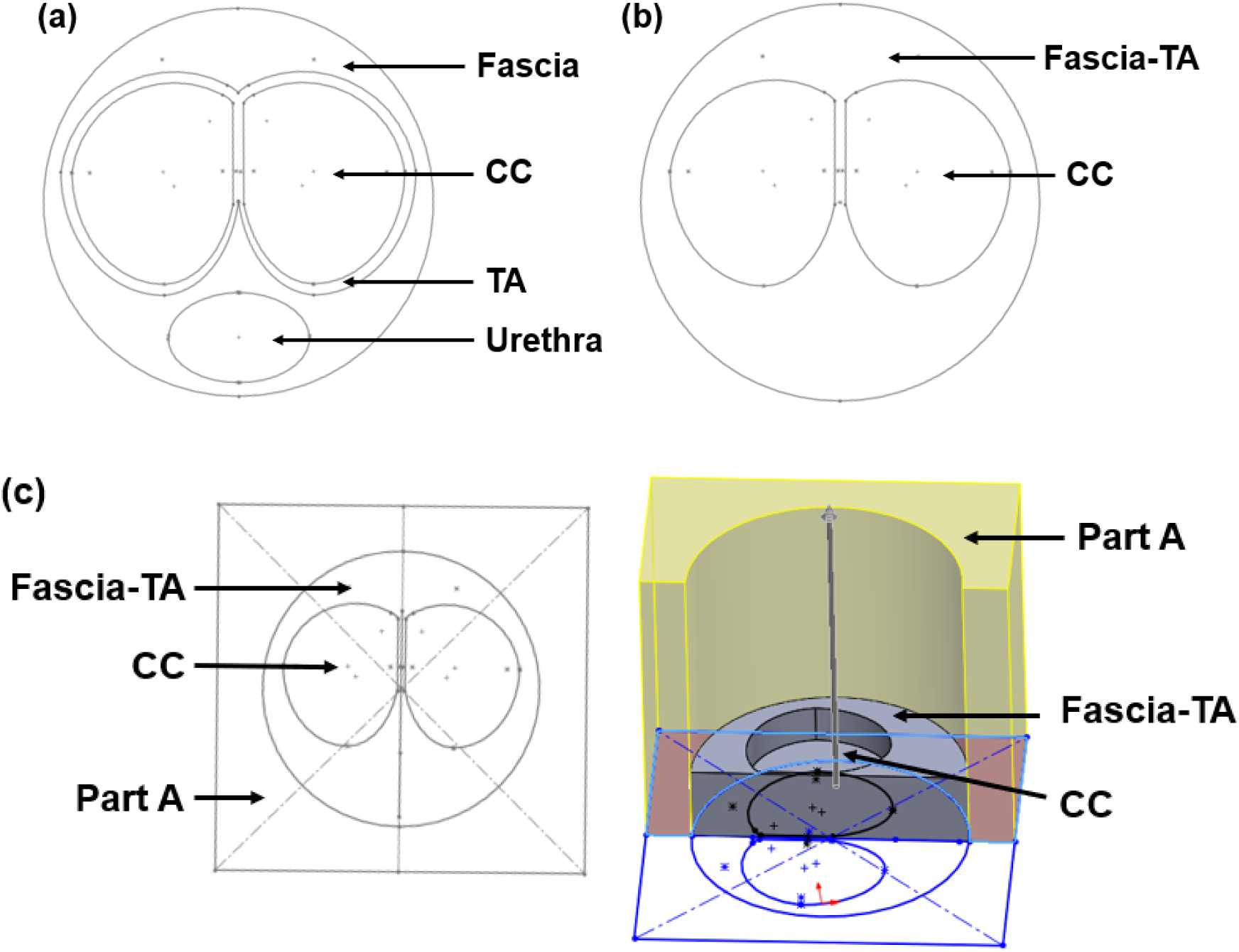
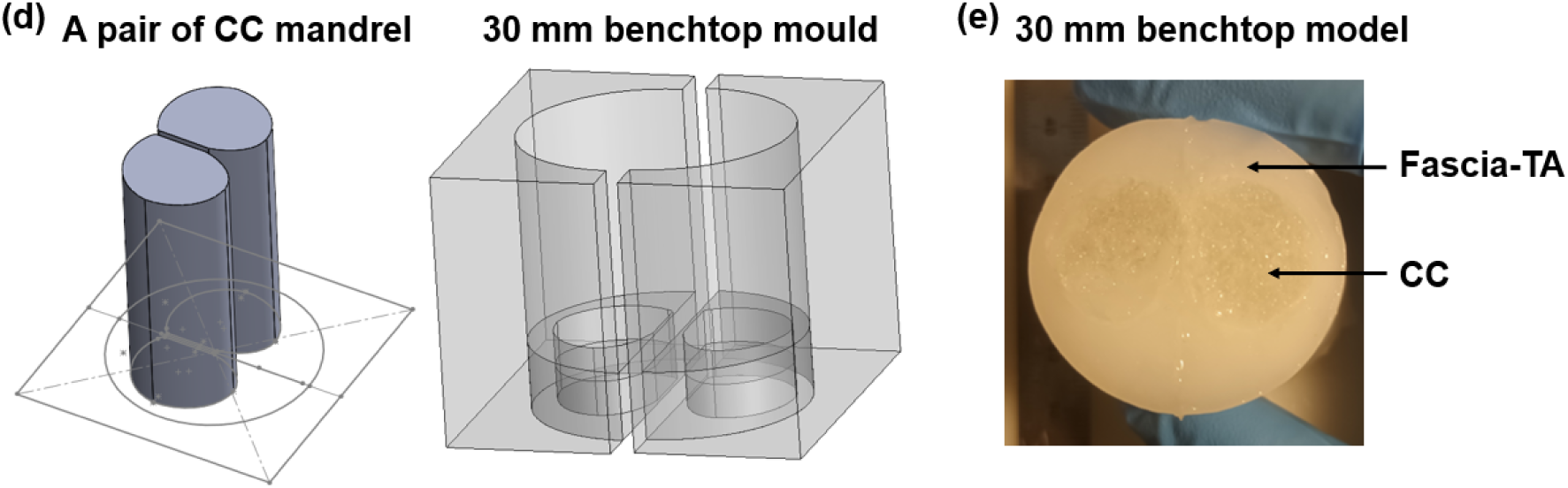
(a) Multi-layered penis anatomy, (b) Simplified penis anatomy for development of benchtop model, (c) Creation of the mould by extruding ‘Part A’ to obtain a 30 mm segment, (d) The geometry of the moulds, (e) The final 30 mm benchtop model.

To develop the fascia-TA layers, the mould was filled with 10% PVA solution, which underwent 2 FTCs. After the end of 1 FTC, the mandrels were removed from CC and it was filled with the porous PVA solution and was allowed to undergo 1 FTC. As mentioned earlier, this solution consisted of 35% milled water-soluble PVA as sacrificial material in a 5% PVA solution. Following the FTCs, the model was placed in DI water to allow particulate leaching of the milled PVA for 72 h. The final benchtop model is shown in Fig. 2e and was stored in DI water. Good adhesion was observed between the fascial-TA layer and CC layer.

### 2.5. Mechanical testing

All the mechanical tests (n=7 for each case) were conducted in a universal testing machine (Zwick Z005, Zwick GmbH & Co. Ulm, Germany) equipped with a 5 N load cell. Compression was performed in a water bath filled with DI water to maintain hydration in the hydrogels. The PVA specimens and human CC had specimen dimensions of 6 ± 0.5 mm (length) and 4.5 ± 0.5 mm (diameter) which were obtained using biopsy punches and were compressed to 50% strain at 10 mm/min.

### 2.6. Inflation testing and ultrasound imaging

Three 30 mm segments from human penises and three individual 30 mm benchtop models were utilised for the IPP inflation testing. The inflation test was conducted using a Boston Scientific AMS 700 CX IPP device, comprising two cylinders (12 cm length, 12 mm diameter), a pump, and a reservoir connected via tubes (Fig. 3a). A digital barometer was affixed to the tube to monitor pressure, maintaining a closed system. Visual assessment of cylinder inflation within the samples at each pressure increment was facilitated using an ultrasound machine -SIEMENS ACUSON S2000™-equipped with a 9L4 linear probe operating at 8 MHz frequency. IPP cylinders were inserted into the corpus cavernosa of the human tissue segments or the porous PVA in the benchtop model, and both were submerged in a water bath (PBS, 37℃), while the barometer, pump, and reservoir remained outside the bath. The ultrasound probe was positioned above the water bath, just outside the water level to visualise the sample, see Fig. 3c. The closed system underwent pressurisation through the pump, while ultrasound images were captured at each pressure increment for subsequent postprocessing.

**Fig. 3:**
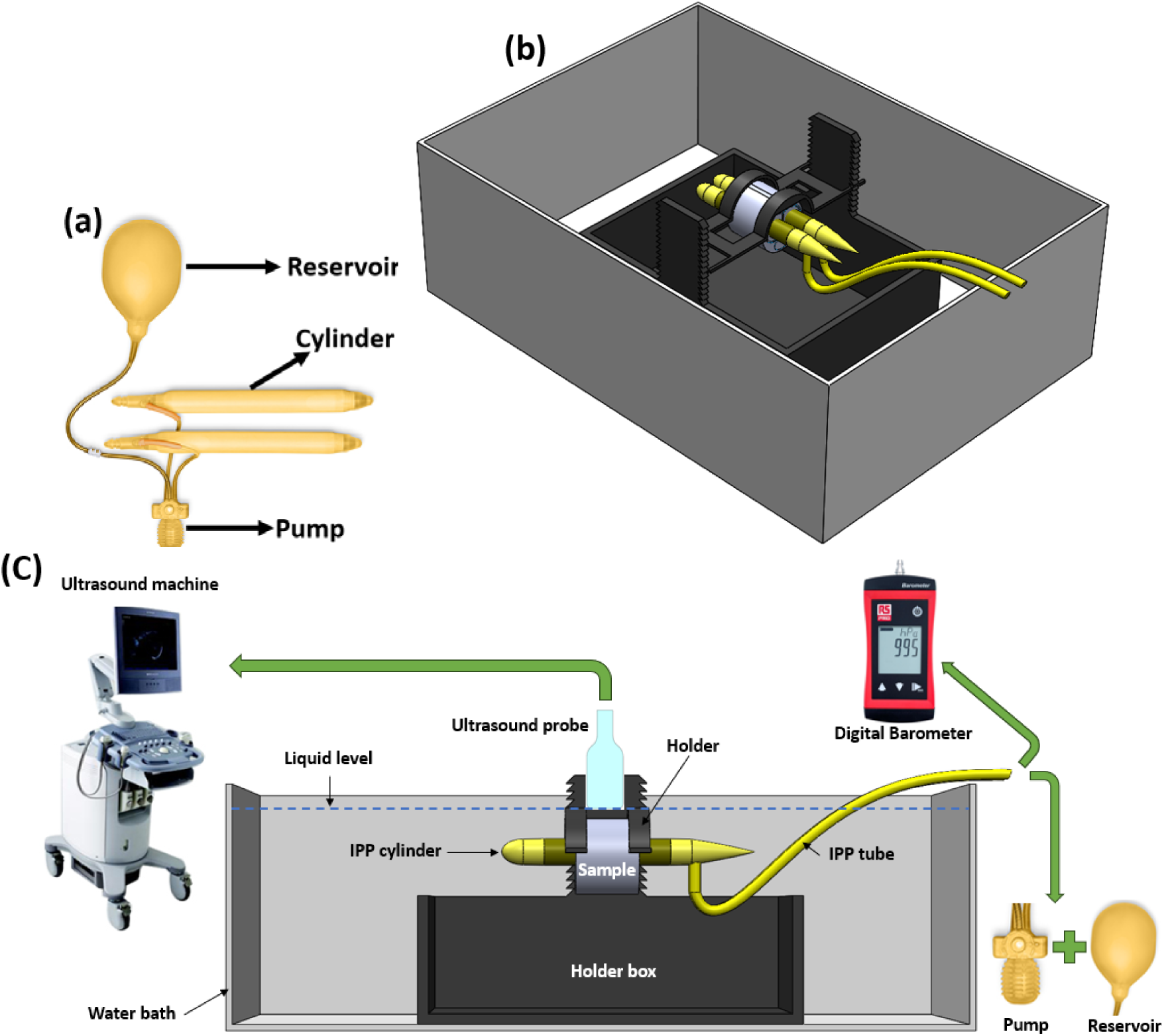
(a) IPP device, (b) isometric schematic view of IPP inflation test set-up, and (c) side view schematic of the test set-up.

To quantify cylinder inflation during the test, the cylinder diameter was measured at each pressure increment. Ultrasound images were imported into ImageJ software (Java 1.8.0_345), and the coordinate points corresponding to the IPP cylinder were exported to enable fitting the best circle through these points using MATLAB (MathWorks, Inc., R2022a), see supplementary materials for further details.

### 2.7. Development of FE model

The cross-section of the penis segments were reconstructed in Abaqus software (Simulia 2022) using the 2D images of the human tissue segments, for example see Fig. 1d. These cross-sections were extruded to create 3D geometries of the segments. Three human penis segments (tissue numbered 1, 2, and 3) and a benchtop model segment were modelled in the software. A 12 mm diameter hole was created in CC for IPP cylinder insertion. Assuming the longitudinal symmetry of the 30 mm human penile segments and bilateral symmetry of the cross-section (see Fig. 1d), a one-fourth model was reconstructed for the finite element (FE) modelling, with corresponding symmetry boundary conditions defined in each plane of symmetry (see Fig. 4b). A pressure of *P*=137.89 kPa (20 psi) was applied to the internal face of the IPP cylinder, based on information from Boston Scientific documents which specify 20 psi as the maximum operating pressure.

**Fig. 4:**
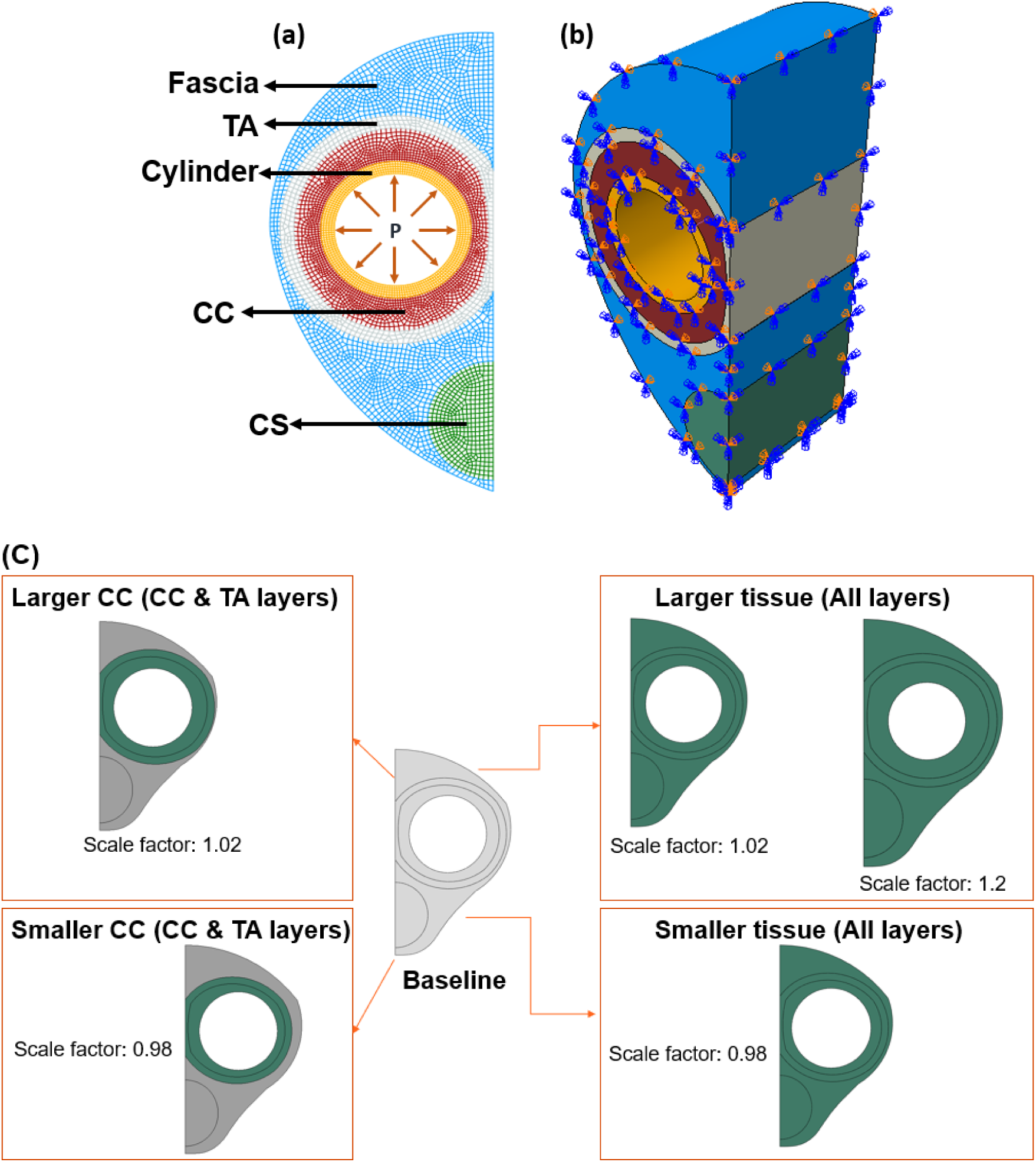
(a) The penile tissue layers (tissue number 2) and IPP cylinder defined in the FE model, (b) the symmetry boundary conditions defined in the 3D FE model. (c) Altering the tissue size in the FE model (tissue number 1).

The penile tissue was presumed to consist of four individual layers: TA, CC, CS, and a fascia layer (see Fig. 4). Hyperelastic material models were employed to define the properties of each tissue layer: the HGO model for TA, the 2^nd^-order Ogden model for CS and CC, and the neo-Hookean model for the fascia layer. The tissue parameters used in this study were adopted from our previous research on human penile tissue characterisation [35], see Table 1. The material properties of the IPP cylinder were obtained through an inverse FE simulation to replicate the pressure-diameter experimental data provided in Boston Scientific documents. The experimental setup involved measuring the cylinder diameter at multiple locations at inflation pressures of 0, 5, 10, 15, and 20 psi using a contact potentiometer, with the cylinder pressurised by water and controlled using a compressed air source and a regulator, see Fig. S3 in supplementary. In the inverse FE approach employed in this study to estimate the cylinder material parameters, the cylinder geometry was created in Abaqus/CAE with outer diameter of 12 mm and a thickness of 1.07 mm, with an internal pressure of 137.89 kPa applied. Due to the highly nonlinear response of the cylinder under inflation, the 2^nd^-order Ogden hyperelastic model was defined. Isight Design Gateway 2022 (Simulia) linked to Abaqus/CAE 2022 was utilised for the inverse FE process. This process aimed to align the simulation pressure-diameter responses with the experimental data by iteratively adjusting the material parameters in the FE model (refer to supplementary document for further details).

**Table 1:** Human penile tissue properties [35].

| Tissue Layer | Material Parameters |  |  |  |  |  |
| --- | --- | --- | --- | --- | --- | --- |
| TA (HGO model) | $C_{10}=100$ kPa | $k_1=1500$ kPa | $k_2=0.1$ | $\kappa=0$ | $D=1e-5$ kPa<sup>-1</sup> | |
| CC (Ogden model) | $\mu_1=0.01$ kPa | $\mu_2=0.012$ kPa | $\alpha_1=20$ | $\alpha_2=20$ | $D_1=1.8$ kPa<sup>-1</sup> | $D_2=0$ |
| CS (Ogden model) | $\mu_1=0.3$ kPa | $\mu_2=0.3$ kPa | $\alpha_1=6$ | $\alpha_2=6$ | $D_1=0.1$ kPa<sup>-1</sup> | $D_2=0$ |
| Fascia (neo-Hookean model) | $C_{10}=0.025$ kPa | $D=0.8$ kPa<sup>-1</sup> | | | | |

To investigate the effects of tissue size variations on inflation test simulations, we altered the cross-sectional dimensions of the 30 mm segment (tissue number 1) using various scale factors: 0.98 (smaller), 1.02 (slightly larger), 1.2 (larger). In addition, we also changed the size of the CC layer (and consequently, the TA layer) with scale factors of 0.98 (smaller CC) and 1.02 (larger CC), while the outer boundary, CS layer size, and TA layer thickness were maintained, without change, as illustrated in Fig. 4c. Additionally, to evaluate the effects of tissue properties, we increased the stiffness of each layer individually (twice the stiffness) by increasing the initial shear moduli in the constitutive models: C_1O_ for the neo-Hookean and HGO models, and µ = µ_1_ + µ_2_ in the Ogden models. As it is hypothesised (and later verified in this paper) that the CC layer is the most influential layer in IPP inflation, a_1_ and a_2_ in the Ogden models were increased up to 24 to examine the effect of increased material nonlinearity.

A mesh study was conducted to determine the required mesh density for a converged solution. The resulting mesh densities were as follows: 103,094 hexahedral elements for human tissue number 1, 120,802 hexahedral elements for human tissue number 2, 114,018 hexahedral elements for human tissue number 3, 147,478 hexahedral elements for the benchtop model, and 63,375 hexahedral elements for the cylinder.

### 2.8. Statistical analysis

The data in this study is presented as mean ± standard deviation. An independent t-test was performed using SPSS 27, IBM, New York, U.S to predict the significant difference between two groups of data. **p<0.01 was considered statistically significant.

## 3. Results

### 3.1. Matching the material properties of human CC to porous PVA hydrogels

The stress-strain curve of the human CC and porous PVA hydrogels under the compression test are plotted in Fig. 5a. The porous PVA specimens show a good matching to that of the material properties of human CC. The initial and final moduli are observed in Fig. 5b. The initial modulus was extracted as the slope of the stress-strain curve in the strain region of 0 – 10% while for final modulus it was estimated in 40%– 50% strain level. Although the final moduli (strains larger than 40%) show a difference between the human CC and the porous PVA, Fig. 5a demonstrates that the porous PVA is able to replicate the stress-strain curve of human CC, and the experimental data for both cases are consistent (see section 4.1).

**Fig. 5:**
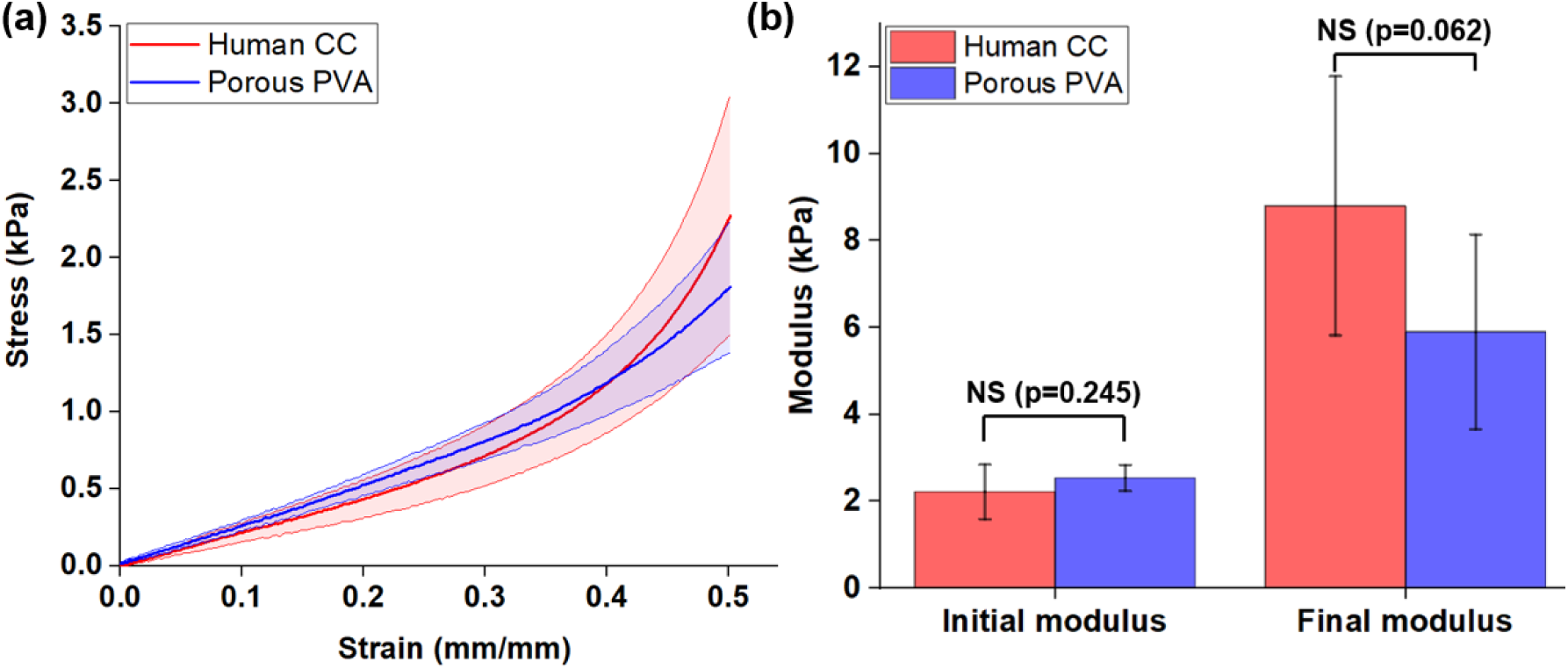
(a) Stress-strain graphs of human CC and porous PVA specimens where the solid lines are the average values and the shaded areas represent standard deviations, (b) Comparison of the initial and final modulus (NS: denotes non-significant difference).

### 3.2. Inflation test

#### 3.2.1. Estimation of IPP cylinder properties

The optimal material parameters estimated for the IPP cylinder are presented in Table 2. These parameters successfully reproduce the inflation test conducted on the cylinder. The comparison of simulation and experimental results are demonstrated in Fig. 6a.

**Fig. 6:**
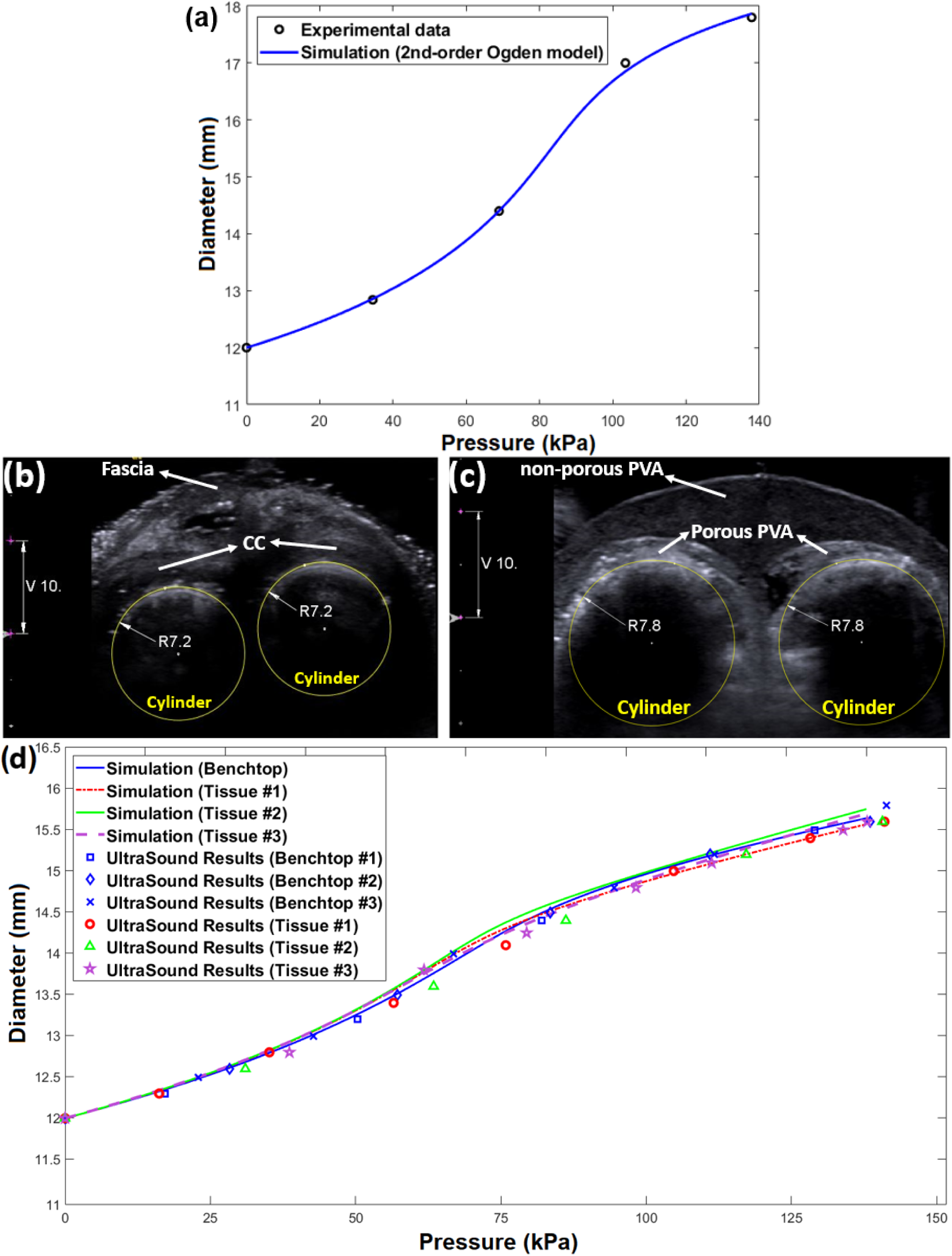
(a) Comparison of the FE simulation of the IPP cylinder expansion under pressure using the material properties shown in Table 2 with the experimental data. Ultrasound images (b) for human tissue segment and (c) benchtop model (all dimensions are in mm). (d) Pressure-diameter experimental data comparing experimental and FE simulation results.

**Table 2:**
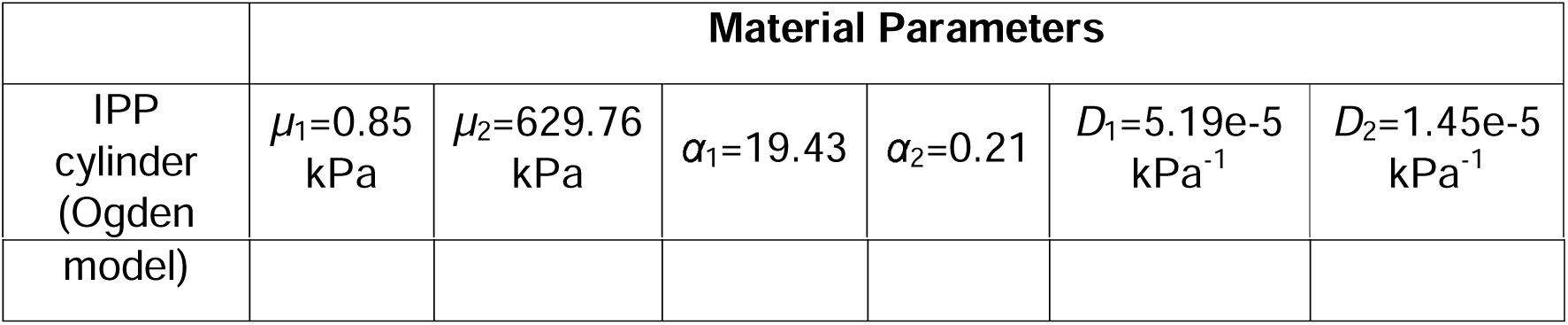
The material properties of IPP cylinder estimated by inverse FE.

|  | Material Parameters |  |  |  |  |  |
| --- | --- | --- | --- | --- | --- | --- |
| IPP cylinder (Ogden) | $\mu_1=0.85$ kPa | $\mu_2=629.76$ kPa | $\alpha_1=19.43$ | $\alpha_2=0.21$ | $D_1=5.19e-5$ kPa<sup>-1</sup> | $D_2=1.45e-5$ kPa<sup>-1</sup> |
| model) |  |  |  |  |  |  |

#### 3.2.2. Model validation

To ascertain the fidelity of our benchtop model, the results obtained from IPP inflation tests on benchtop model segments are compared to human tissue segments, supported by FE simulations. These results reveal a robust concurrence between the responses, as depicted in Fig. 6d. The pressure-diameter curves exhibited by the benchtop models, made of a combination of porous and non-porous PVA, closely replicate those of human tissues. The FE simulations also show a good agreement with the experimental data, see Fig. 6d.

In addition to comparing pressure-diameter results, it is important to examine the tissue deformations caused by IPP cylinder inflation by comparing the principal strain contours from simulations (at *P*=137.89 kPa), as shown in Fig. 7. This comparison indicates that the strain levels and distributions are consistent and comparable in all cases, with some discrepancies due to geometric differences, such as the maximum principal strain at the sharp corner of the fascia layer where it interfaces with the TA layer. The contours are even more consistent in the CC layers, where the principal strain ranges are approximately between -0.5 and -0.73, with the largest strains occurring at the outer boundary. This consistency validates the FE model developed in this study for simulating IPP inflation within human penile tissue. The absolute value of maximum principal strains (the principal strains with the largest absolute magnitude) is used to visualise both positive and negative principal strains.

**Fig. 7:**
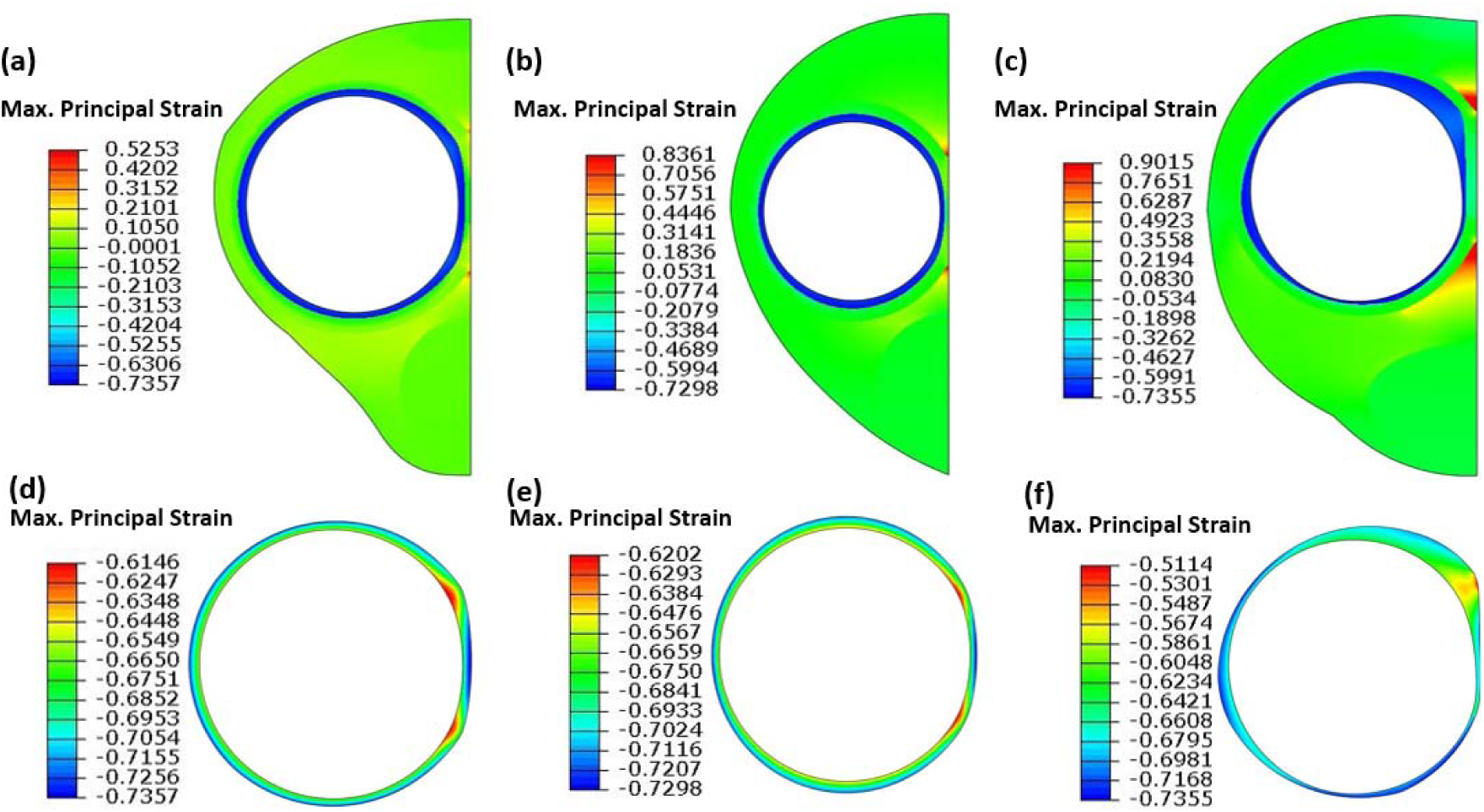
Absolute maximum principal nominal strains calculated by FE simulations (at maximum applied pressure); (a) tissue number 1, (b) tissue number 2, (c) tissue number 3, (d) the CC layer of tissue number 1, (e) the CC layer of tissue number 2, and (f) the CC layer of tissue number 3.

#### 3.2.3. Effects of size and properties

Focusing on the effects of tissue size, it is observed that the results are identical for different dimensions until approximately *P*=70 kPa (10.15 psi), as shown in Fig. 8a. After this pressure, tissue dimension becomes a factor, with larger dimensions resulting in increased cylinder inflation. Notably, slight changes in all tissue dimensions yield similar outcomes to changes in CC with the same scale factor (highlighted curves in Fig. 8a). This highlights the strong influence of the CC layer size in the IPP inflation test.

Examining the effect of tissue layer properties (Fig. 8b) indicates that increasing the stiffness of fascia and CS layers has negligible impact on inflation results, even with a tenfold increase in stiffness (see Fig. S5 in the supplementary). The stiffness of CC has the most significant influence on tissue inflation, deviating from the baseline curve at pressures around 65 kPa (9.42 psi). In contrast, the curve representing stiffer TA maintains the same results as the baseline until approximately 90 kPa (≈ 13 psi), with less deviation compared to the stiffer CC curve.

**Fig. 8:**
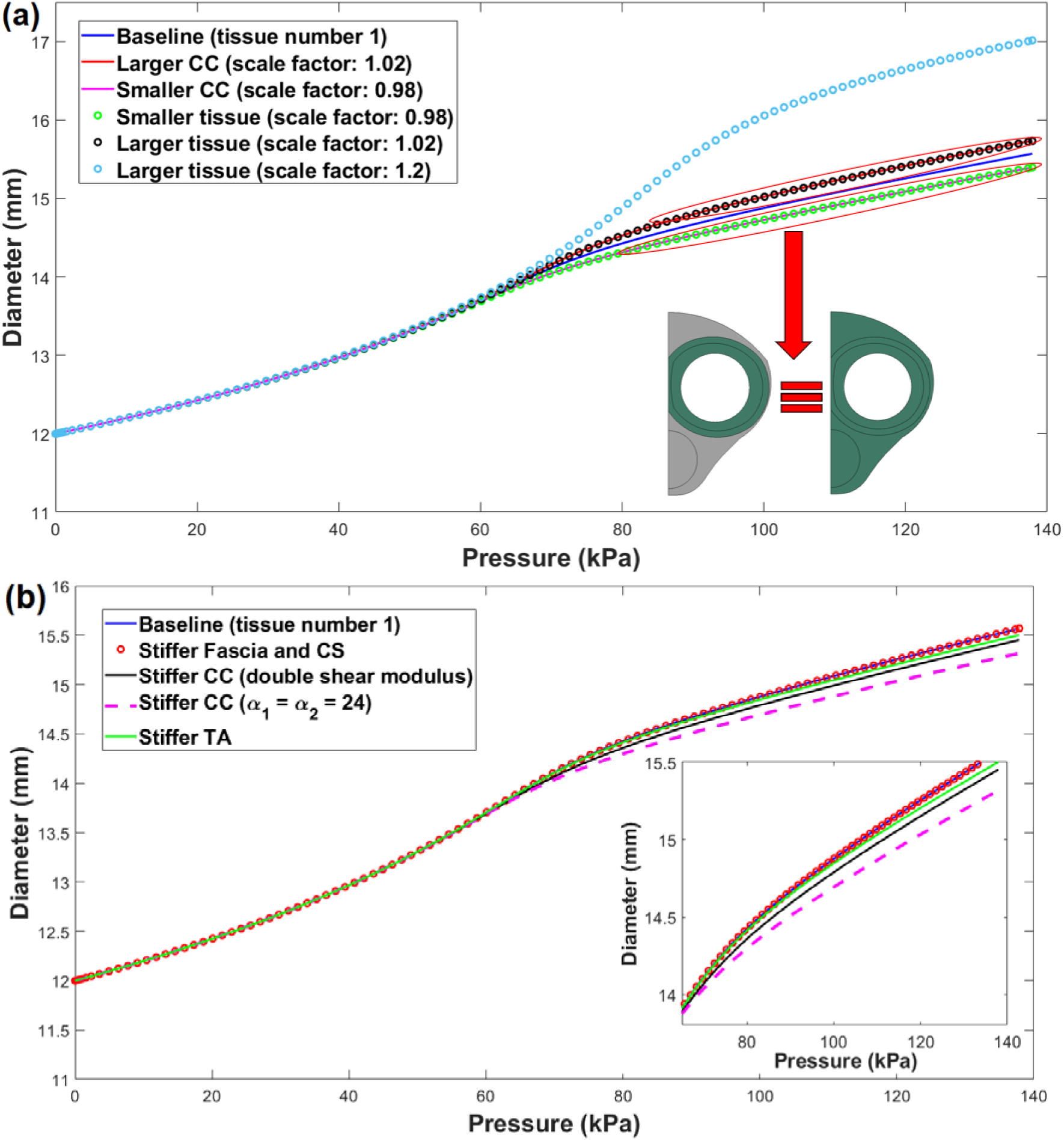
Pressure-diameter FE simulation results to investigate tissue layer (a) sizes and (b) properties.

## 4. Discussion

### 4.1. Validity of CC representation using porous PVA

Our simulation and experimental results support the suitability of porous PVA hydrogels as a surrogate for human CC tissue. As demonstrated in Fig. 5a, the stress-strain curve of porous PVA aligns well with that of human CC up to moderate strain levels. A discrepancy appears in the final modulus at strains beyond 40% and it is also noteworthy that the maximum compressive strain induced in the CC during the IPP inflation can reach values up to 0.73 (73%) (as shown in Fig. 7), which is higher than the 0.5 (50%) strain used in compression tests on human CC [35] (Fig. 5a). While further experimental data at higher strains would be ideal, it is shown that the porous PVA prepared in this study as a representative of CC layer is able to replicate the mechanical behaviour of the human CC under IPP inflation, as evidenced by the experimental pressure-diameter data shown in Fig. 6d. In addition, the simulation results confirm that modest increases in CC stiffness have only a minimal effect on IPP inflation behaviour (see section 4.2), as shown in Fig. 8b. Overall, the benchtop model reliably mimics the behaviour of human penile tissue under IPP inflation, despite slight differences between the mechanical properties of human CC and porous PVA.

### 4.2. Effects of size and properties

Figs. 8a and 8b, along with supplementary Fig. S5, clearly demonstrate the dominant role of the CC layer in governing the tissue’s mechanical response during IPP inflation. The inflation test simulations for various tissue layer sizes and properties show that the inflation test is insensitive to both size and mechanical properties at low IPP pressures, as demonstrated in Figs. 8a and 8b. At higher pressures, although the fascia and CS layers continue to have minimal influence, the CC and TA layers become more significant, with the CC layer clearly playing the dominant role in the IPP inflation scenario. However, the results for a slightly stiffer CC, shown in Fig. 8b, exhibit only minor deviations from the baseline, with maximum relative errors of just 0.7% and 1.6% for a doubled initial shear modulus and an increased nonlinear parameter a, respectively. While ten times stiffer CC and TA demonstrate much greater effect on IPP inflation, see Fig. S5 in the supplementary. The maximum relative error percentage is calculated using 100 x (D_base_ - D_stiff_)⁄D_base_, where D_base_ and D_stiff_ are the maximum cylinder diameters at the maximum pressure *P*=137.89 kPa for the baseline and stiffer CC curves, respectively.

It is important to note that extreme sizes were not utilised due to geometric constraints; very small CC sizes would not accommodate the 12 mm IPP cylinder, while increasing CC size with higher scale factors would exceed tissue boundaries. Additionally, less stiff tissue layer properties were not considered for brevity, as they would increase IPP inflation (cylinder diameter). Furthermore, while there are additional parameters (as shown in Table 1) that can influence the hyperelastic behaviour of each tissue layer, this study focused on increasing initial shear moduli and α parameters (in CC) for simplicity. Tissues defined as twice the stiffness were chosen to maintain reasonable stiffness for penile tissues [35] encompassing tissue diversity or diseases such as Peyronie’s disease, which stiffen the TA [36], [37]. However, parametric results demonstrating tissues ten times stiffer are presented in Fig. S5 in the supplementary.

### 4.3. Limitations

One limitation of this study is that the ultrasound is only able to visualise the upper half of the samples’ cross-section, as can be seen in Figs. 8a and 8b, due to the size of the segments. However, the IPP cylinders into the samples remain approximately circular after inflation as confirmed by FE simulations. Therefore, it is possible to measure the radius of cylinders using the arc shown in the ultrasound results. Refer to the supplementary file for more information about cylinder size measurements.

In addition, conducting compression tests at higher strain levels (up to 0.7-0.75) would be advantageous for validating tissue parameters over a wider range of strains. Damage and viscoelastic analyses on human CC will be complementary to provide a more comprehensive understanding of the impact of IPP inflation on the tissue.

### 4.4. Conclusion

A benchtop model has been successfully developed with a spongy CC layer using porous PVA that showed mechanical properties consistent with those of human CC. Following this, a 30 mm segment penile benchtop model was created using different concentrations and FTCs of PVA, with the outer fascial layers made of non-porous PVA. The suitability of the benchtop model was validated by inflating it and comparing the results to those of human tissue and an FE model informed by human tissue tests. The inflation test results revealed a strong correlation between these models, indicating that both the benchtop and FE models can be used as pre-clinical testbeds in the future.

## Acknowledgements/Funding sources

This publication has emanated from research conducted with the financial support of Science Foundation Ireland (SFI) under grant number 12/RC/2278_2 and Boston Scientific Limited (Clonmel).

