## Supplementary Text and figures for "Urological benchtop and *in silico* models validated by human penile tissue inflation tests"

1. **Diameter measurement**

To measure the cylinder diameters in the IPP inflation test, the image captured from the ultrasound machine for each pressure increment were imported to ImageJ software (Java 1.8.0_345). The points on the outer boundary of cylinders were manually selected in ImageJ and extracted as text files to represent (X, Y) data points in a cartesian coordinate system. These coordinate points were called in a MATLAB (MathWorks, Inc., R2022a) code to fit the best circle through these points using *circfit* function. For instance, the following figures illustrate the circle fitted through the data points for human tissue number 1.

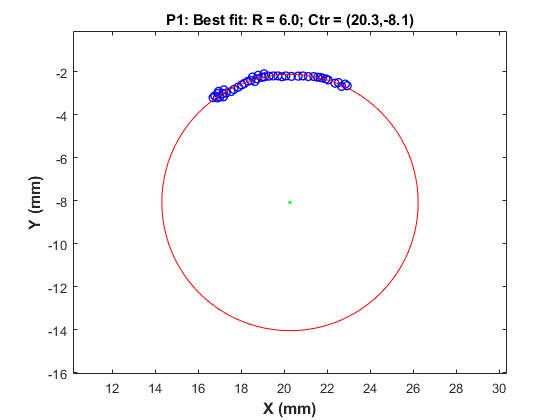

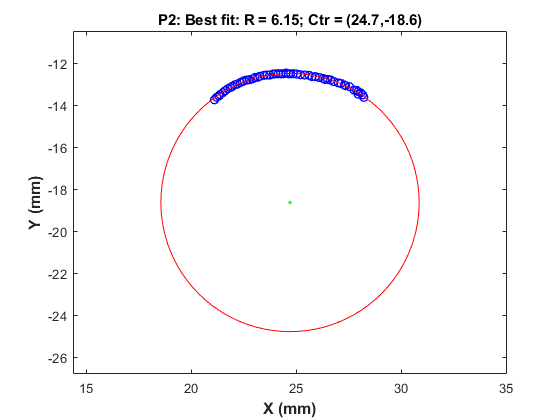

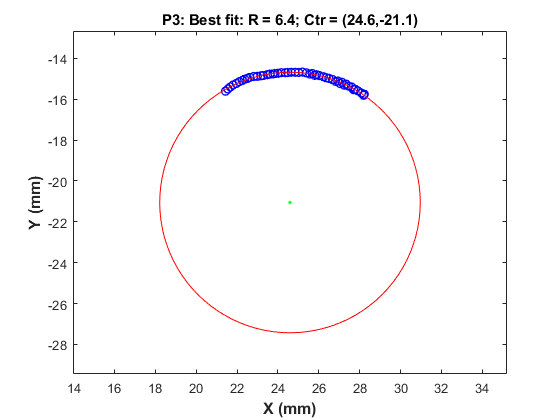

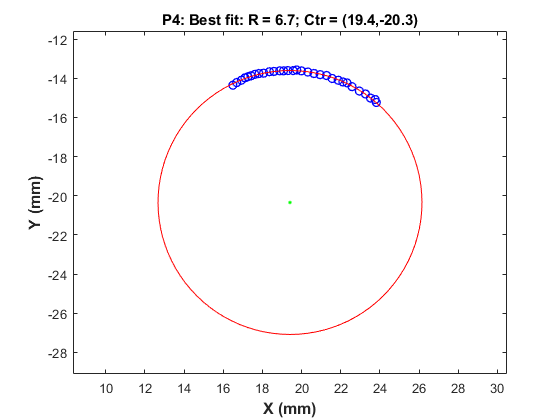

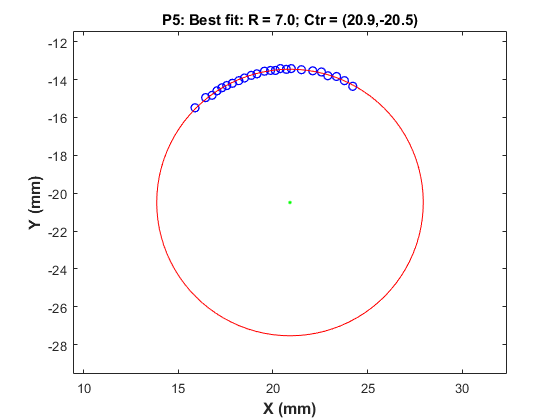

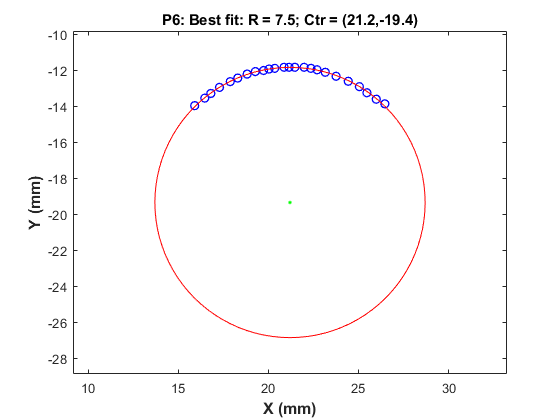

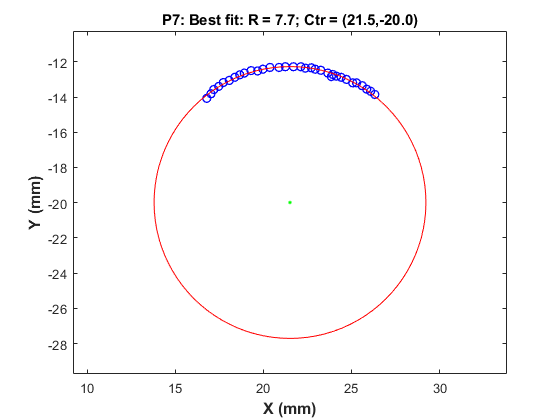

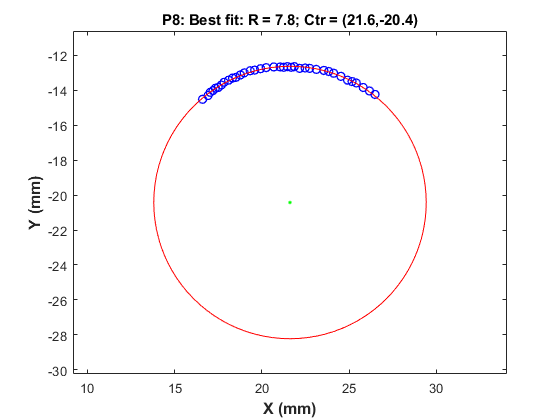

**Fig. S1:** The best circle fitted through data points to measure the IPP cylinder size during the inflation test, in which *R* and *Ctr*, respectively, represent circle radius and centre point, and *P* denotes the absolute pressure value at each increment (*P*_1_=0, *P*_2_=16.20 kPa, *P*_3_=35.16 kPa, *P*_4_=56.53 kPa, *P*_5_=75.84 kPa, *P*_6_=104.73 kPa, *P*_7_=128.24 kPa, *P*_8_=140.99 kPa).

2. **Inverse FE approach**

In the inverse FE approach used in this study to estimate the material parameters of IPP cylinder, the .cae and .odb files of the FE model (Abaqus 2022, Simulia) was imported to the Isight Design Gateway 2022 (Simulia) to read the material models, initial guesses, and values for pressure and displacement. The desired outputs are transferred to a Calculator unit to set the simulation results (here, pressure and displacement) for each time step as arrays. The simulation results were compared to the experimental data in the Data Matching unit, as shown in Fig. S2a. The sum of the absolute difference between points on the simulation and experiments was defined as the objective function $\varphi$, as follows:

$$\varphi=\sum_{i}^{N} \left| \chi_{i}^{exp}-\chi_{i}^{sim} \right|$$

(1)

where $\chi_{i}^{exp}$ and $\chi_{i}^{sim}$, represent experimental and simulated values, respectively. This inverse FE approach uses the Hooke-Jeeves algorithm as a direct search optimisation technique to minimise the objective function. In this algorithm, the relative step size, step size reduction factor, and termination step size are 0.02, 0.5, 1.0e-6, respectively. The inverse FE flowchart is shown in Fig. S2b.

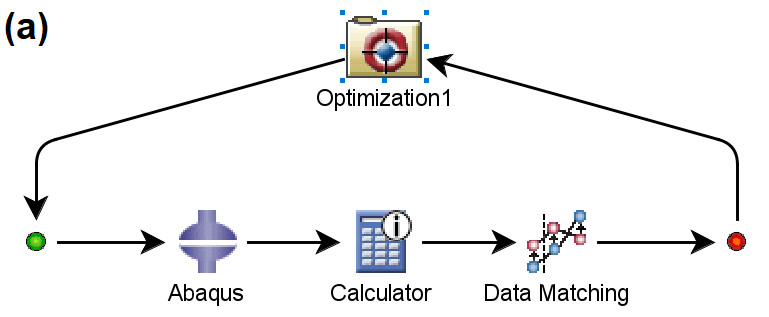

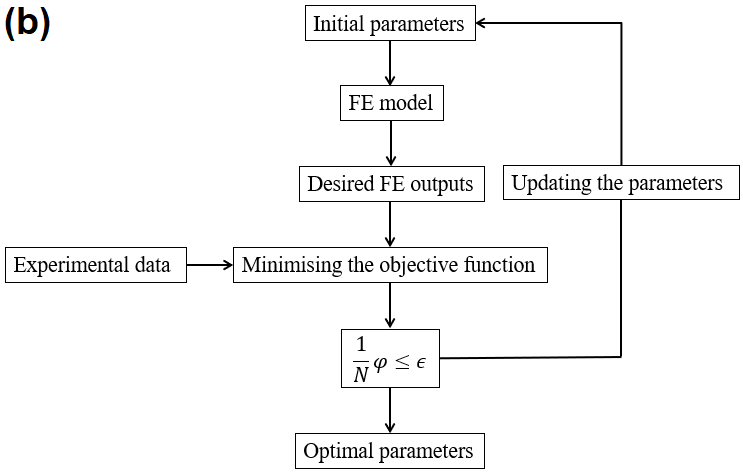

**Fig. S2:** (a) the inverse FE approach used in Isight software, (b) the inverse FE flowchart, where the termination criterion, *ϵ*, is set to 0.05, and *N* demotes the number of experimental data points.

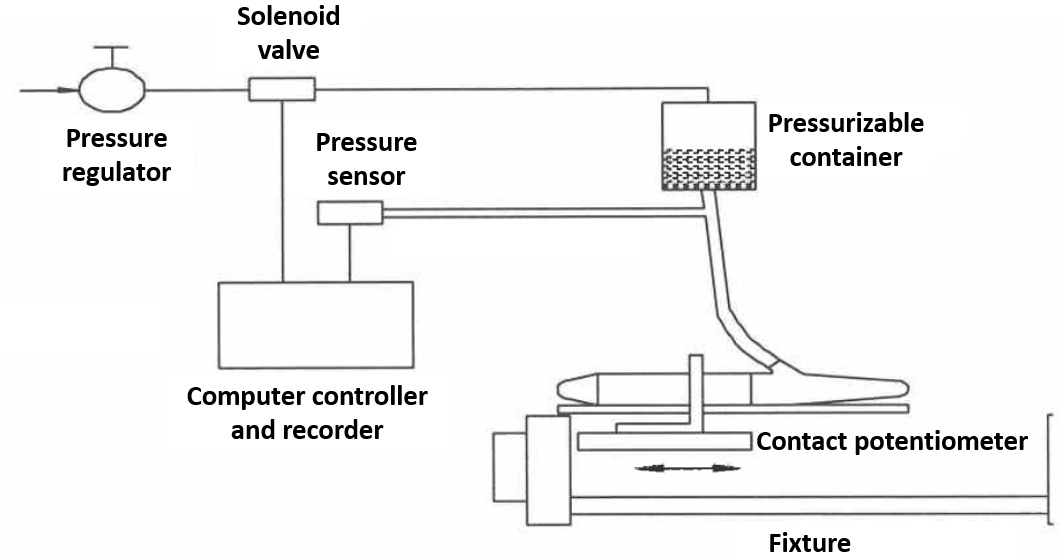

**Fig. S3:** Test set-up to measure cylinder diameter and length (internal Boston Scientific Corporation document)

3. **Tissue properties**

Fig. S4 illustrates the absolute maximum principal nominal strains of CC layers calculated by FE model using stiffer material properties.

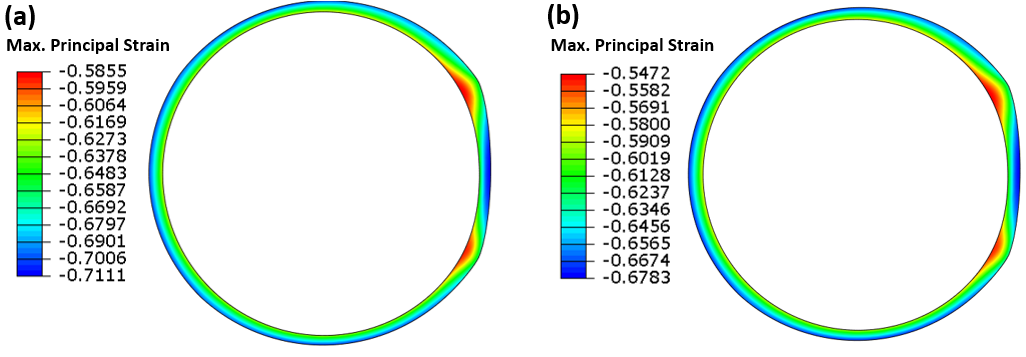

**Fig. S4:** Absolute maximum principal nominal strains of CC layers calculated by FE simulations (at maximum applied pressure); (a) *µ*_1_=0.02 kPa and *µ*_2_=0.024 kPa, and (b) *α*_1_=*α*_2_=24. The remaining parameters are the same as those in Table 1.

The IPP inflation simulations indicate that the fascia and CS layers have minimal impact on the inflation results, even with a tenfold increase in stiffness, as illustrated in Fig. S5. Note that a tenfold increase in stiffness refers to multiplying the initial shear modulus of the layer by ten.

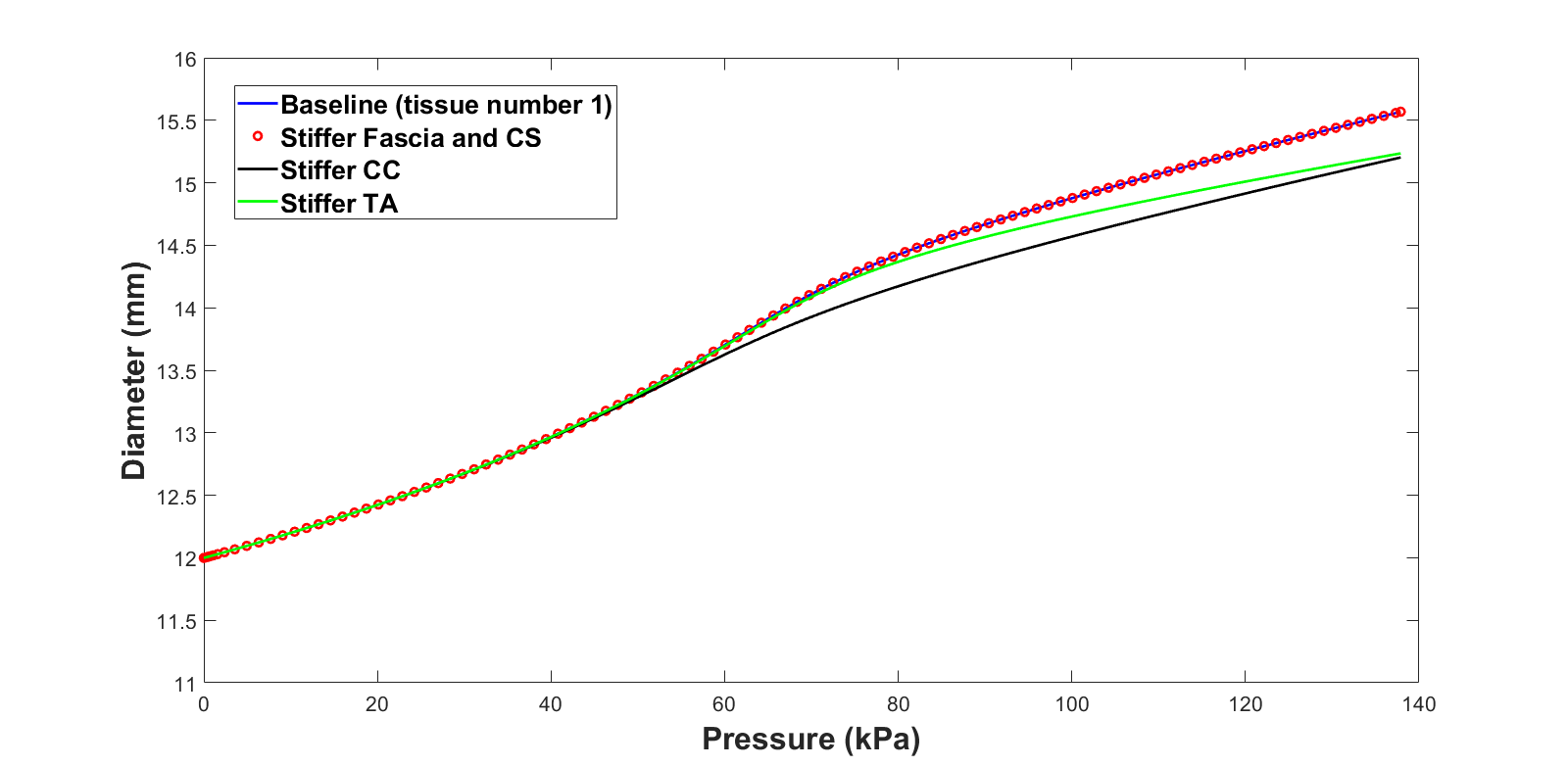

**Fig. S5:** Pressure-diameter FE simulation results for tenfold stiffer tissue properties.

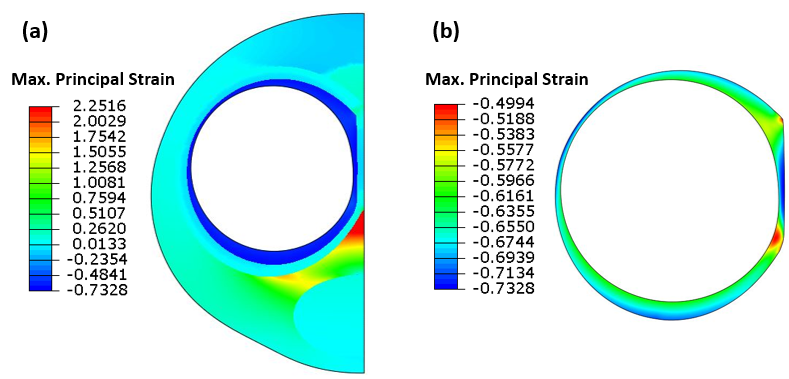

**Fig. S6:** Absolute maximum principal nominal strains calculated by FE simulations (at maximum applied pressure); (a) benchtop model, (b) the CC layer of benchtop model.
